# A multiple genome alignment workflow shows the impact of repeat masking and parameter tuning on alignment of functional regions in plants

**DOI:** 10.1101/2021.06.01.446647

**Authors:** Yaoyao Wu, Lynn Johnson, Baoxing Song, Cinta Romay, Michelle Stitzer, Adam Siepel, Edward Buckler, Armin Scheben

**Affiliations:** Institute for Genomic Diversity, Cornell University, Ithaca, NY USA 14853; Agricultural Genomics Institute at Shenzhen, Chinese Academy of Agricultural Sciences, Shenzhen, China; U.S. Department of Agriculture-Agricultural Research Service, Ithaca, NY USA 14853; Simons Center for Quantitative Biology, Cold Spring Harbor Laboratory, Cold Spring Harbor, NY; Department of Molecular Biology and Genetics, Cornell University, Ithaca, NY USA 14853

**Author notes:** These authors contributed equally (Yaoyao Wu, Lynn Johnson).

## Abstract

Alignments of multiple genomes are a cornerstone of comparative genomics, but generating these alignments remains technically challenging and often impractical. We developed the *msa_pipeline* workflow (https://bitbucket.org/bucklerlab/msa_pipeline) based on the LAST aligner to allow practical and sensitive multiple alignment of diverged plant genomes with minimal user inputs. Our workflow only requires a set of genomes in FASTA format as input. The workflow outputs multiple alignments in MAF format, and includes utilities to help calculate genome-wide conservation scores. As high repeat content and genomic divergence are substantial challenges in plant genome alignment, we also explored the impact of different masking approaches and alignment parameters using genome assemblies of 33 grass species. Compared to conventional masking with RepeatMasker, a *k*-mer masking approach increased the alignment rate of CDS and non-coding functional regions by 25% and 14% respectively. We further found that default alignment parameters generally perform well, but parameter tuning can increase the alignment rate for non-coding functional regions by over 52% compared to default LAST settings. Finally, by increasing alignment sensitivity from the default baseline, parameter tuning can increase the number of non-coding sites that can be scored for conservation by over 76%.

## Introduction

Multiple sequence alignment is a key challenge in comparative genomics and evolutionary studies (Chowdhury and Garai 2017; Armstrong et al. 2019). As the number of novel genomes being generated is rapidly accelerating, researchers rely on robust tools that can scale from dozens to hundreds of genomes. Many tools are available for pairwise or multiple alignment of genome sequences (Frith and Kawaguchi 2015; Marçais et al. 2018; Armstrong et al. 2020; Minkin and Medvedev 2020). However, these tools generally require a range of inputs such as a phylogenetic tree and repeat masking information. Pairwise alignment tools such as LASTZ and LAST also need their outputs to be post-processed before subjecting them to multiple alignment using a different tool. In addition, many tools do not scale well to large sets of plant genomes. The many requirements and types of software involved can make the seemingly straightforward task of multiple sequence alignment technically challenging for individual researchers. In this work, we therefore developed the practical *msa_pipeline* to generate multiple sequence alignments from a reference genome and a set of query genomes. The *msa_pipeline* relies on the LAST aligner and aims to minimize the amount of user effort required to rapidly produce a high-quality multiple alignment. We tested the computational efficiency of the pipeline and the impact of a range of repeat masking and alignment parameters using public grass genome sequences. Overall, we present the publicly available *msa_pipeline* and recommend repeat masking and alignment strategies that enhance alignment of genic and intergenic regions of diverged plant genomes.

### Features and implementation of *msa_pipeline*

The *msa_pipeline* only requires a set of masked genomes in FASTA format as input, outputting a multiple sequence alignment in MAF format (Figure 1). Dependencies are handled using Docker/Singularity and snakemake is deployed as a workflow manager. We used the LAST alignment tool for pairwise alignment, rather than the faster minimap2, because the high sensitivity of LAST (Frith and Noé 2014) makes it more suitable for comparison of diverged genomes. High sensitivity is important for many downstream analyses of the alignment, because it facilitates alignment of functional sequences such as promoters and enhancers that are located in more variable intergenic regions.

**Figure 1.**
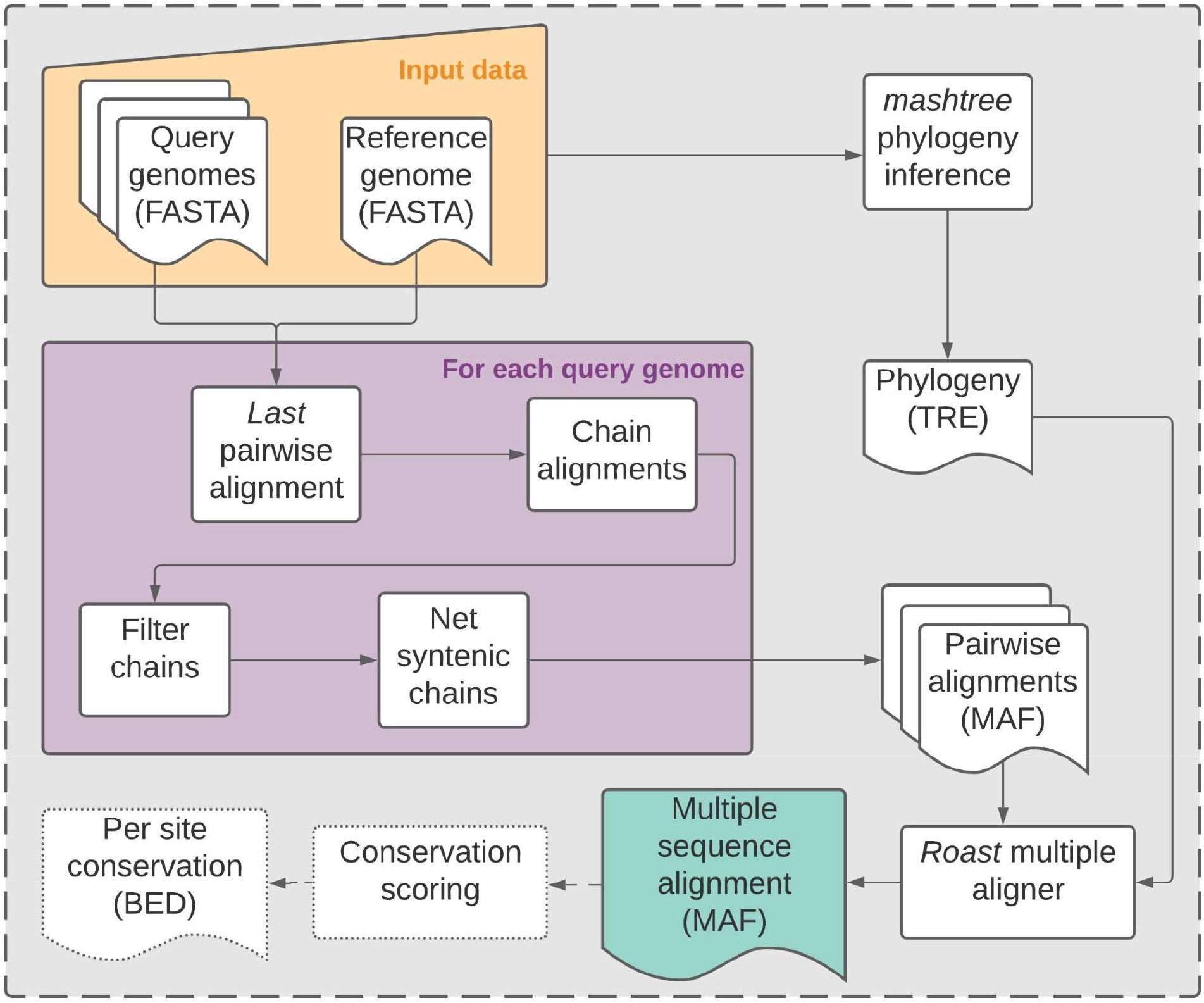
Schema describing the snakemake multiple sequence alignment pipeline

Pairwise alignment to the reference genome is conducted in parallel, with the main pipeline bottleneck being multiple sequence alignment using the single-threaded ROAST program. The pipeline outputs multiple alignments in MAF format. We provide scripts to use the alignment to generate genome-wide conservation scores calculated with GERP++ (Davydov et al. 2010), phastCons or phyloP (Siepel et al. 2005). The runtime and memory usage of *msa_pipeline* shows its efficiency compared to the powerful but resource-intensive Cactus aligner (Table 1).

**Table 1.** Computational resources used by *msa_pipeline* for multiple alignment using different species sets and masking approaches. A comparison estimated resource requirements for the Cactus multiple aligner highlights the relative speed and low memory use of the *msa_pipeline*.

| Clade | # Species | Masking | *msa_pipeline* |  | Cactus 1.2.3* |  |
| --- | --- | --- | --- | --- | --- | --- |
|  |  |  | Runtime with 12 threads (h) | Max memory (Gb) | Estimated runtime with 12 threads (h) | Estimated max memory (Gb) |
| PACMAD | 14 | soft | 31.8 | 15.8 | ~2400 | ~850 |
| PACMAD | 14 | hard | 27.8 | 16.8 | - | - |
| BOP | 19 | soft | 51.0 | 22.1 | - | - |
| BOP | 19 | hard | 45.4 | 22.7 | - | - |

\*Due to its high resource requirements, the Cactus aligner resource use estimate is based on an alignment prepared for a separate study using 10 PACMAD species (including three shared with the *msa\_pipeline* set). The alignment was conducted with 96 threads and runtime per thread is conservatively estimated by dividing total runtime by the thread count. Runtime was rounded down to the nearest hundred hours and memory was rounded down to the nearest 50 Gb.

### Benchmarking and improving multiple alignment in plant genomes

#### Selecting alignment benchmarking metrics

Measuring the accuracy of alignments between distantly related species is challenging because ground-truth alignments are generally unknown. Studies have therefore measured alignment accuracy by focusing on partial alignments of conserved functional sequences such as exons (Sharma and Hiller 2017; Frith, Hamada, and Horton 2010) or by relying on simulated sequences (Armstrong et al. 2020). To reduce biases caused by simulation parameters or by an exclusive focus on coding sequence, we measured accuracy based on alignments of functional sequences in coding and non-coding regions. Specifically, we calculated precision, recall and F_1_ score (harmonic mean of precision and recall) of functional regions, assuming that alignments of non-functional regions were false positives (see Methods for further details). Although this simplifying assumption is unlikely to generally be the case, the resulting approximate measures are useful for benchmarking alignment quality in the functional regions of the genome that are most important for the majority of downstream analyses.

### Appropriate repeat masking can improve multiple alignment performance

A major obstacle to accurate and efficient alignment is the large proportion of repetitive sequence found in most plant genomes. In contrast to masking tools like RepeatMasker that rely on repeat databases, approaches such as RED (Girgis 2015) or KMER (Song et al. 2020) try to avoid database bias by using repetitive *k*-mers (nucleotide sequences of *k* length) in the genome to identify repeats. Here, we compared RepeatMasker, RED and KMER and tested their impact on subsequent multiple sequence alignment in grasses. We selected species from the PACMAD grass clade (subfamilies Panicoideae, Aristidoideae, Chloridoideae, Micrairoideae, Arundinoideae, and Danthonioideae) which diverged ∼32.4 mya (Cotton et al. 2015) as well as species from the BOP grass clade (subfamilies Bambusoideae, Oryzoideae, and Pooideae) which diverged ∼80 mya (Christin et al. 2014). We found substantial differences between all three masking methods, impacting the amount of putative false positive masking in coding, open chromatin regions and non-coding functional regions (see Methods for definition of these regions).

In maize, compared to KMER, RepeatMasker masked an additional 28.89% of CDS and 38.96% of non-coding functional regions (Figure 2A, Table S1). KMER also masked substantially less sequence than RED (Figure 2A and Table S1). Overall, KMER displayed the most favorable trade-off between the masking rate and the rate of masked coding and non-coding functional sequence across most genomes (Figure 2B, Figure S1B, Table S2; see Supplementary Results and Supplementary Data). KMER, however, failed to mask substantial numbers of repeats in fragmented genome assemblies such as those of *Dichanthelium oligosanthes* and *Eragrostis tef*.

**Figure 2.**
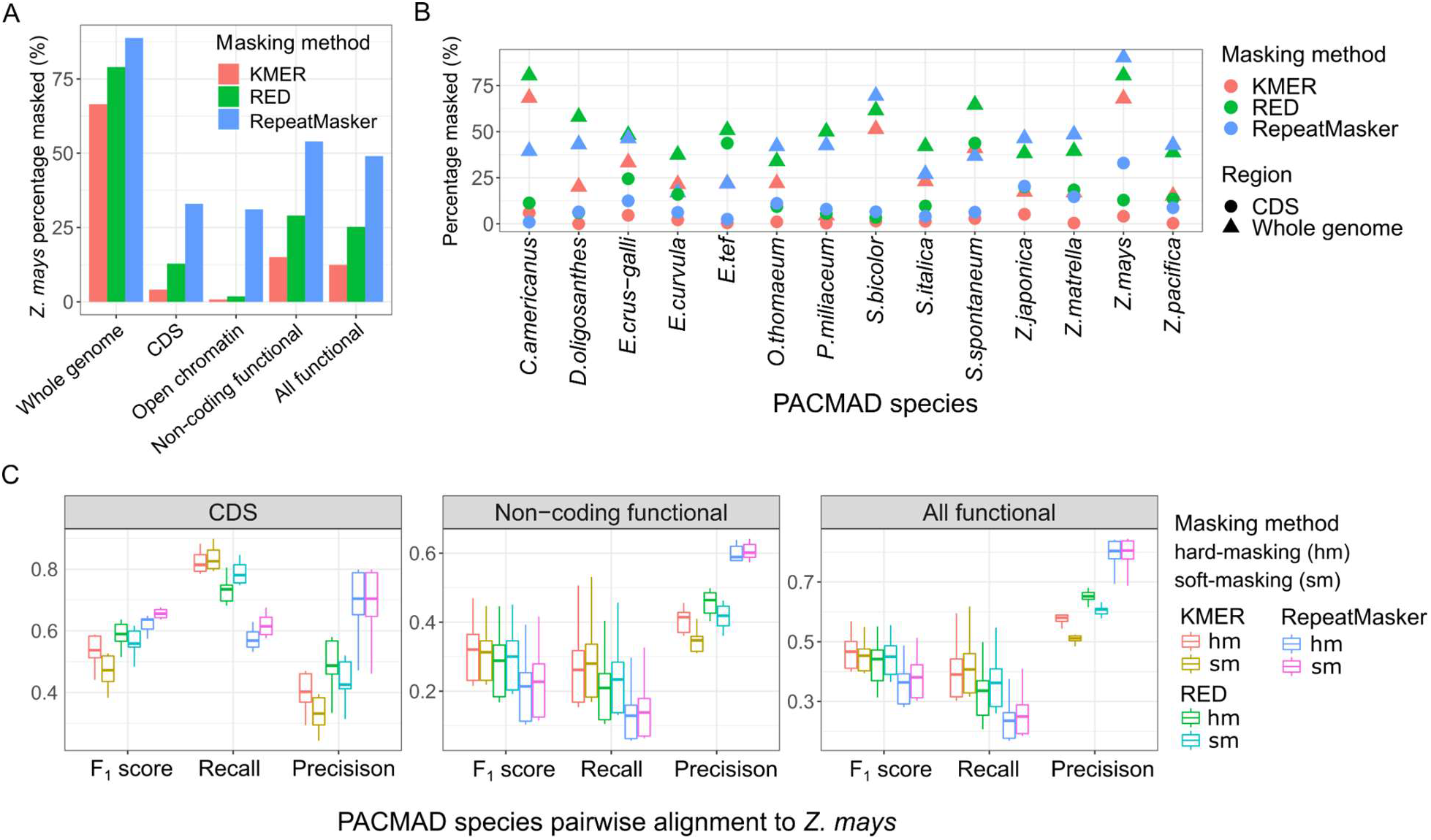
Impact of repeat masking methods on alignment of functional genomic regions. (A) The masking rate for different genomic regions in maize using three masking methods. (B) The masking rate for the whole genome and for CDS in 14 species of the PACMAD clade. (C) Boxplots of pairwise alignment performance (see Methods) of 13 species of the PACMAD clade aligned to maize.

Based on analysis in the PACMAD clade, genomes masked with KMER produced sensitive alignments (mean F_1_= 0.4670 for pairwise alignment; multiple alignment F_1_= 0.4809) with higher alignment rates of functional sequence than those masked with RepeatMasker (mean F_1_= 0.3569 for pairwise alignment; multiple alignment F_1_= 0.3686) and those masked with RED (mean F_1_= 0.4284 for pairwise alignment; multiple alignment F_1_= 0.4506) (Figure 2C, Figure S1C and S1D; see Supplementary Data). Our results suggest that using *k*-mer-based masking improves alignment, with hard-masking performing comparably to soft-masking while also providing minor improvements in runtime (Table 1 and Table S3).

### Exploration of alignment parameter space shows potential for improving intergenic alignment rates

Alignment parameters such as substitution matrices and gap penalties can have a substantial effect on alignment (Frith, Hamada, and Horton 2010). Often default alignment settings are based on testing in mammalian genomes that are less repetitive and diverse than those of many plants. To explore the alignment parameter space for grass genomes, we tested 750 different combinations of ten LAST parameters including gap penalties and substitution matrices for multiple alignments (Table S4). By approximating recall and precision as measures of alignment performance, we assessed 750 multiple alignments of a 5Mb and 1.4 Mb syntenic region in the grass clades known as PACMAD and BOP (Figures S2-S5). Although we found that some of the best alignments were generated using default LAST alignment parameters (recall = 0.2823, precision = 0.9095, F_1_ = 0.4309) and Cactus alignment (recall = 0.4040, precision = 0.8478, F_1_ = 0.5472), alternative LAST parameter combinations showed substantial differences including some improvements in alignment performance compared to the default parameters (Table S5, Table S6 and Table S7). Across the 750 parameter combinations, coding regions (recall = 0.48-0.78; precision = 0.47-0.99) showed substantially higher recall than non-coding regions (recall = 0.02-0.39; precision = 0.63- 0.93, Table S5).

The default LAST penalty matrix and parameters favor precision over recall, which we found leads to low alignment rates in intergenic regions for divergent genomes like those in the PACMAD grass clade. In this study, we selected the parameter combination ‘LAST strict’ (Table S6) with equal precision compared to LAST default parameters but a recall of 0.35, corresponding to a 23% increase from the default (Table S7). This gain in recall is mainly attributable to use of the HOXD70 penalty matrix and lower penalization of gaps (Table S6). The parameter combination ‘LAST relaxed’ (Table S6) further decreases the gap existence cost (parameter *-a*), elevating the recall to 0.57 while maintaining a precision over 0.85. This parameter combination produces an alignment with similar precision and recall but substantially lower computational cost compared to the Cactus alignment in both the PACMAD and BOP clade (Figure 3, Figure S5, and Table S7).

**Figure 3.**
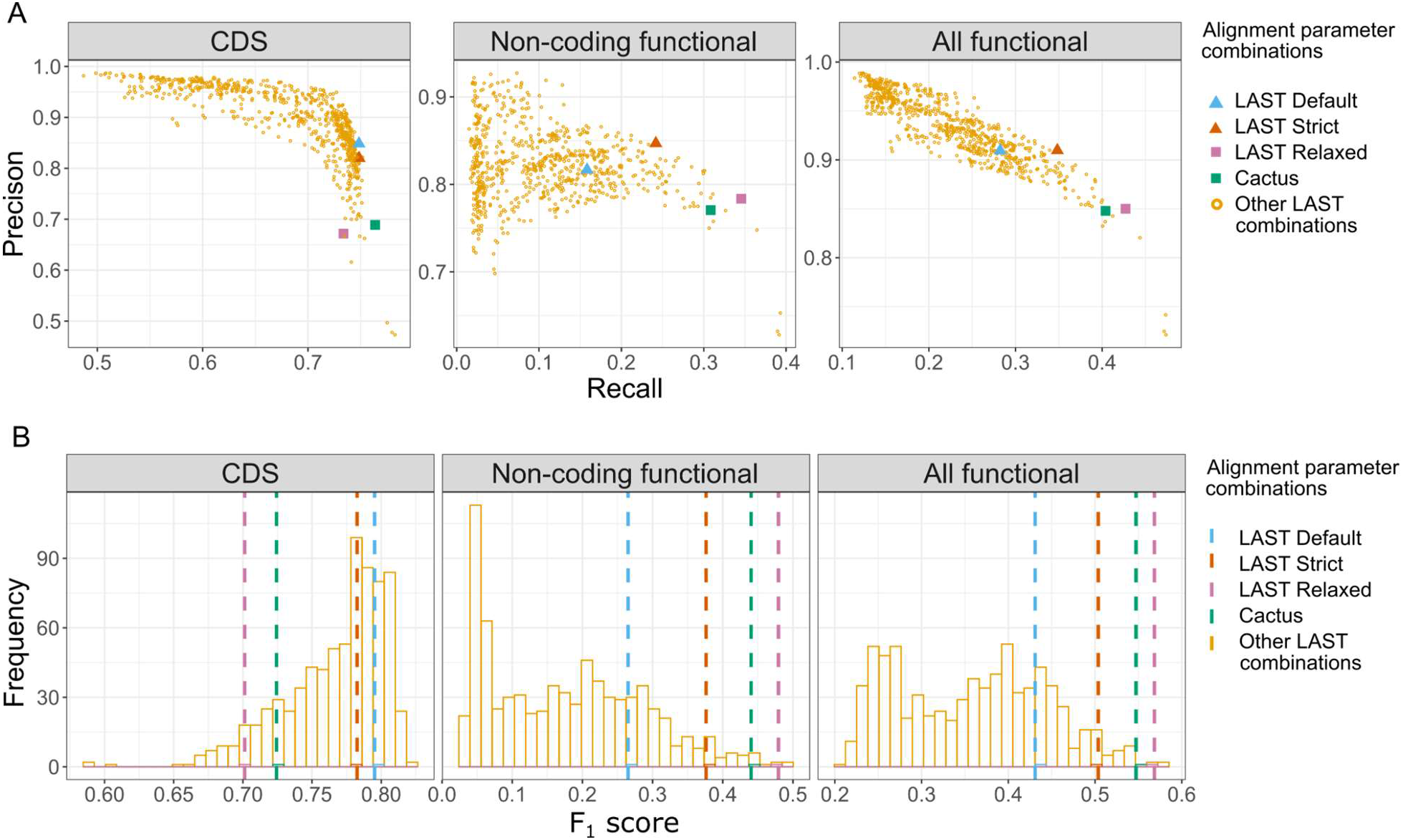
Multiple alignment performance of 750 LAST parameter combinations in different genomic regions in the PACMAD grass clade. Tested parameter combinations are compared to the alignment performance of default LAST parameters and the Cactus 1.2.3 aligner based on (A) recall and precision as well as the (B) F_1_ score.

### Multiple alignment parametrization facilitates detection of genomic conservation

To evaluate how much the multiple alignment affects estimates of genomic conservation, we calculated the GERP conservation score based on the previously introduced 750 alignments of PACMAD and BOP generated with different alignment parameter combinations. In PACMAD, the number of sites that had sufficient alignment depth (>=3 species) to produce a conservation score ranged from 92,437 to 3,843,983 (Figure 4A), and the number of detected conserved sites ranged from 16,559 to 131,820 (Figure 4B; see Supplementary Data). The LAST default parametrization led to detection of 98,193 conserved sites. The ‘LAST strict’ parametrization led to detection of 113,253 conserved sites, corresponding to a 15.35% increase compared to the default (1.61% increase in CDS region, 75.75% increase in non-coding functional region). In line with this result, the parameter combination ‘LAST relaxed’ elevated the number of detected conserved sites by 19.77% (−4.43% in CDS region, 114.31% increase in non-coding functional region) (Figure 4B and Table S8; see Supplementary Data). We found a similar substantial increase in the detectable conserved sites in the BOP clade (Figure S6; see Supplementary Data). The mean Pearson’s correlation (r) in conservation scores between the PACMAD and BOP clades in syntenic regions was moderate (r=0.25) with limited variability between alignment parameter combinations (Figure S7). Taken together, these results suggest that the HOXD70 substitution matrix combined with a relatively low gap-open penalty is preferable to the default LAST substitution matrix and gap-open penalty for detection of plant conserved non-coding elements.

**Figure 4.**
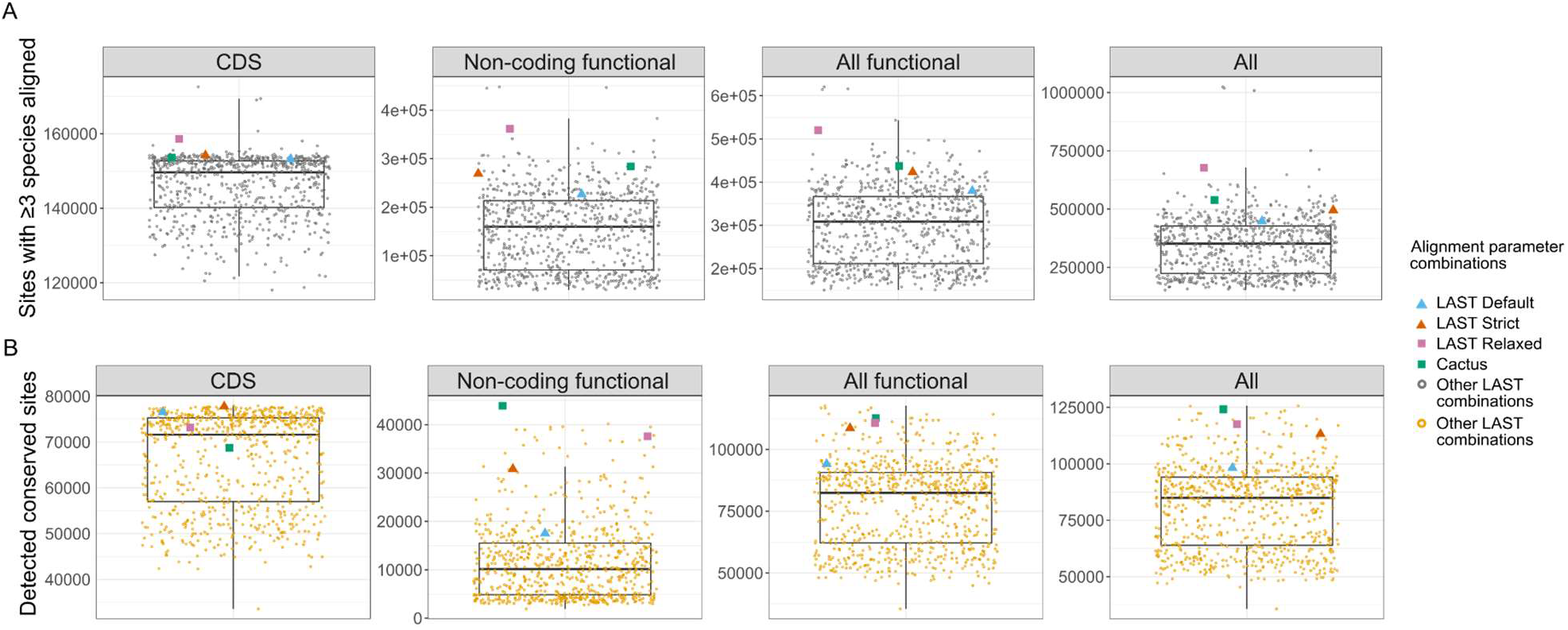
The GERP performance of 750 LAST parameter combinations in different genomic regions in the PACMAD grass clade. (A) The number of sites with sufficient alignment depth (>=3 species) to be scored for conservation in different genomic regions in the PACMAD grass clade. (B) The number of conserved sites in different genomic regions in the PACMAD grass clade.

## Outlook

The *msa_pipeline* leverages existing tools to provide a practical solution for rapid multiple alignment of genomes with minimal user effort. For divergent plant genomes, different repeat masking approaches had limited impact on the alignment rate, but reduction of gap-related alignment penalties boosted alignment rates of non-coding functional elements. We anticipate that the accelerating pace of genome sequencing and assembly will generate rich resources for genome-scale multiple alignments that drive biological discovery in plants.

## Methods

### Repeat masking approaches

Repeats often cannot be aligned accurately between genomes. For this reason, repetitive sequences are often replaced with ‘N’s (hard-masked) or set to lowercase (soft-masked) and treated differently from non-repetitive sequences during alignment. Repeat masking with popular methods such as RepeatMasker generally relies on libraries of repeat elements that are aligned to genomes to identify known repeats. Here, repeat masking was carried out on the genome assemblies using RepeatMasker 4.1.1 with the RepBase 20181026 database of Viridiplantae and a custom set of repeats mined from each genome using RepeatModeler. A drawback is that repeat elements not similar to those in the library will not be masked and, conversely, non-repetitive functional elements with similarities to repeats may be erroneously masked (Bayer, Edwards, and Batley 2018). To compare kmer-based masking approaches to RepeatMasker, we therefore also conducted masking with RED and a novel kmer-based approach (Song et al., 2020) that we refer to as KMER.

### Selection of syntenic regions for alignment analysis

Whole genome alignment is computationally demanding (Table 1). To accelerate comparison of multiple alignments constructed with a range of parameters, we used a subset of genomic sequences from our target species. Specifically, we used MCScan (Wang et al. 2012) to select a syntenic region that is common to grass genomes and contains 100 genes based on the *Sorghum bicolor* GCF_000003195.3 genome (Figure S2; see Supplementary Data). This allowed us to compare alignment results from two distinct clades of grasses known as the BOP and PACMAD clades. *Oryza longistaminata* was excluded from mini-genome analyses due to poor alignment rates (see Supplementary Data). The reference genome used for the 18 selected BOP species was rice (version IRGSP-1.0) and the reference genome for the 14 selected PACMAD species was maize (version B73V4).

### Sampling the alignment parameter space

We performed multiple alignment with the *msa_pipeline* for PACMAD clade species and BOP clade species using three sets of differently masked sequences (RepeatMasker, RED, KMER) for each clade. Each masking approach was furthermore tested with hard-masked and soft-masked sequences. We varied 10 LAST pairwise alignment parameters to explore the parameter space (Table S4), including parameters controlling gap/mismatch penalty sizes, number of initial matches and simple repeat masking. A total of 750 parameter combinations were randomly sampled from the parameter space. Two custom substitution penalty matrices (RETRO and RETRO SIMPLE; see Supplementary Data) were generated based on observed substitution rates in aligned maize retrotransposons. Briefly, we used MAFFT alignments of 5’ and 3’ long terminal repeats (LTRs) of individual retrotransposon copies from Stitzer et al. (2019) to count base substitutions that have accumulated since the TE inserted, using the seg.sites function implemented in ape v5.4 (Paradis and Schliep 2018). This provides an empirical measure of substitution rates in maize, reflecting the high rate of transitions.

### Evaluation of alignments

We focus on the alignment of functional elements of the genome as a measure of alignment quality. We use a broad definition of these functional elements, including non-coding functional regions (promotors, UTRs, introns, open chromatin) and coding regions (CDS).

We define *a* as the number of bases of *Zea mays* functional elements with at least half of the query species aligned, while *e* is defined as the total number of bases of *Zea mays* functional elements.

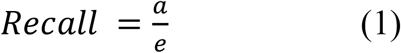

Thus we define approximate alignment recall as shown in equation 1.

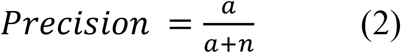

In equation 2, we define the number of aligned non-functional intergenic bases as *n* and use them to help calculate approximate alignment precision. A key assumption here is that intergenic regions distant from genes and with inaccessible chromatin are enriched for erroneous alignments compared to our defined functional regions. This assumption is a caveat for our calculation of precision, because false positives are identified based on this assumption rather than a ground truth.

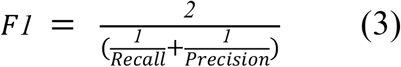

Finally, we can calculate the F_1_ score using our calculations of alignment recall and precision.

### Alignments affect the detection of genomic conservation

To assess how the alignment affects the inference of genomic conservation, we calculated conservation using GERP with the *msa_pipeline* in the PACMAD and BOP clade respectively. For each alignment generated from the 750 parameter combinations, we used a fixed neutral tree and considered all sites with Rejected Substitution (RS) scores greater than 80% of the maximum RS score to be conserved. The threshold for considering a site conserved in BOP was RS=1.568 and the threshold in PACMAD was RS=1.072.

To further explore the site to site alignment, we used Pearson’s correlation of GERP RS scores between the PACMAD and BOP clades. We expect a substantial proportion of conservation to be clade-specific and thus uncorrelated, limiting the maximum correlation possible. However, we cautiously consider an increase in correlation as a potential indicator for improvements in alignment of functional sequences conserved across grass clades.

We used LAST alignment to lift-over genomic coordinates between the rice genome (the reference for BOP) and the maize genome (the reference for PACMAD). For the sites that could be lifted over between rice and maize, we then calculated the correlation of GERP RS scores between PACMAD and BOP across the genome and for different functional genomic regions.

## Code availability

The *msa_pipeline* code is available at https://bitbucket.org/bucklerlab/msa_pipeline/.

## Acknowledgements

This work was supported by NSF (grant IOS-1822330) and USDA-ARS. M.C.S. was supported by NSF PRFB 1907343. The authors acknowledge the Texas Advanced Computing Center (TACC) at The University of Texas at Austin for providing HPC resources that have contributed to the research results reported within this paper. Further computational work was done using resources of the Cornell Biotechnology Resource Center Bioinformatics Facility (Computational Biology Service Unit, CBSU). Jeffrey Ross-Ibarra provided helpful comments throughout the analysis and writing. We also acknowledge the assistance of Ritika Ramani in writing and testing the bash wrapper for the pipeline.

## Supplement

### Supplementary data

The Supplementary Data files S1-S10 are listed below and can be downloaded from FigShare (https://doi.org/10.6084/m9.figshare.14691318.v1).

**Supplementary Data S1**. Repeat masking results for grass genomes using three masking methods

**Supplementary Data S2**. Alignment metrics for whole genome pairwise alignment

**Supplementary Data S3**. Alignment metrics for whole genome multiple alignment

**Supplementary Data S4**. List of genes in the Sorghum mini-genome

**Supplementary Data S5**. List of genes in the Oryza sativa mini-genome

**Supplementary Data S6**. Mini-genome alignment rates across species

**Supplementary Data S7**. Multiple alignment summary metrics for 750 alignment parameters combinations

**Supplementary Data S8**. Alignment performance of high-performing parameters compared to the default parameters and the Cactus aligner

**Supplementary Data S9**. Genomic conservation scoring for multiple alignments generated with 750 alignment parameter combination compared to the default parameters

**Supplementary Data S10**. Custom LAST penalty matrices based on substitution rates calculated from *Z. mays* transposon alignments

## Supplementary results

Compared to KMER, RepeatMasker masked an additional 22.76% (whole genome), 28.89% (CDS), 30.28% (open chromatin) and 38.98% (non-coding functional) in maize (Figure 1A). And when compared to RED, RepeatMasker masked 10.25%, 20.08%, 29.28% and 24.97% more in the whole genome, CDS region, open-chromatin and non-coding functional region (Figure 1A). RepeatMasker thus had the highest masking rate but for many species, it also masked the highest proportion of functional elements occurring in coding and open chromatin sequence, and non-coding functional region (Figure 1A). KMER displayed the most favorable trade-off between the masking rate and the rate of masked coding and open chromatin sequence. This difference between RepeatMasker and KMER was not specific to maize, it was noticeable in most grass genomes (Figure 1B, Figure S1). However, KMER performance declined in cases where the genome was poorly assembled, as is the case for *Dichanthelium oligosanthes* and *Eragrostis tef*.

To further investigate the impact of masking on pairwise alignment, we analyzed the alignment recall, precision and F_1_ score for CDS, open chromatin regions, non-coding functional regions and all functional regions (Figure 1C). Using pairwise alignments of 32 grass species, the average F_1_ scores were 0.40 (KMER), 0.35 (RED), 0.29 (RepeatMasker), and 0.32 (unmasked). This showed that KMER produces the highest F_1_ score and that hard-masked alignment performs similarly to soft-masked alignment but with a significant speed-up (Figure 1, Figure S1, Table 1). For the multiple alignment of all species in the PACMAD clade, the F_1_ is 0.4309 (KMER), 0.3919 (RED), 0.3203 (RepeatMasker), and 0.3542 (unmasked) for functional regions. Although alignments with KMER masking received the highest F_1_ score in both the PACMAD and BOP clade (Figure S1D), the repeat masking approach only had a substantial impact on alignment rates when using hard-masking. These results suggest that the *msa_pipeline* is not sensitive to the repeat-masking approach for soft masked genomes.

## Supplementary figures and tables

**Figure S1.**
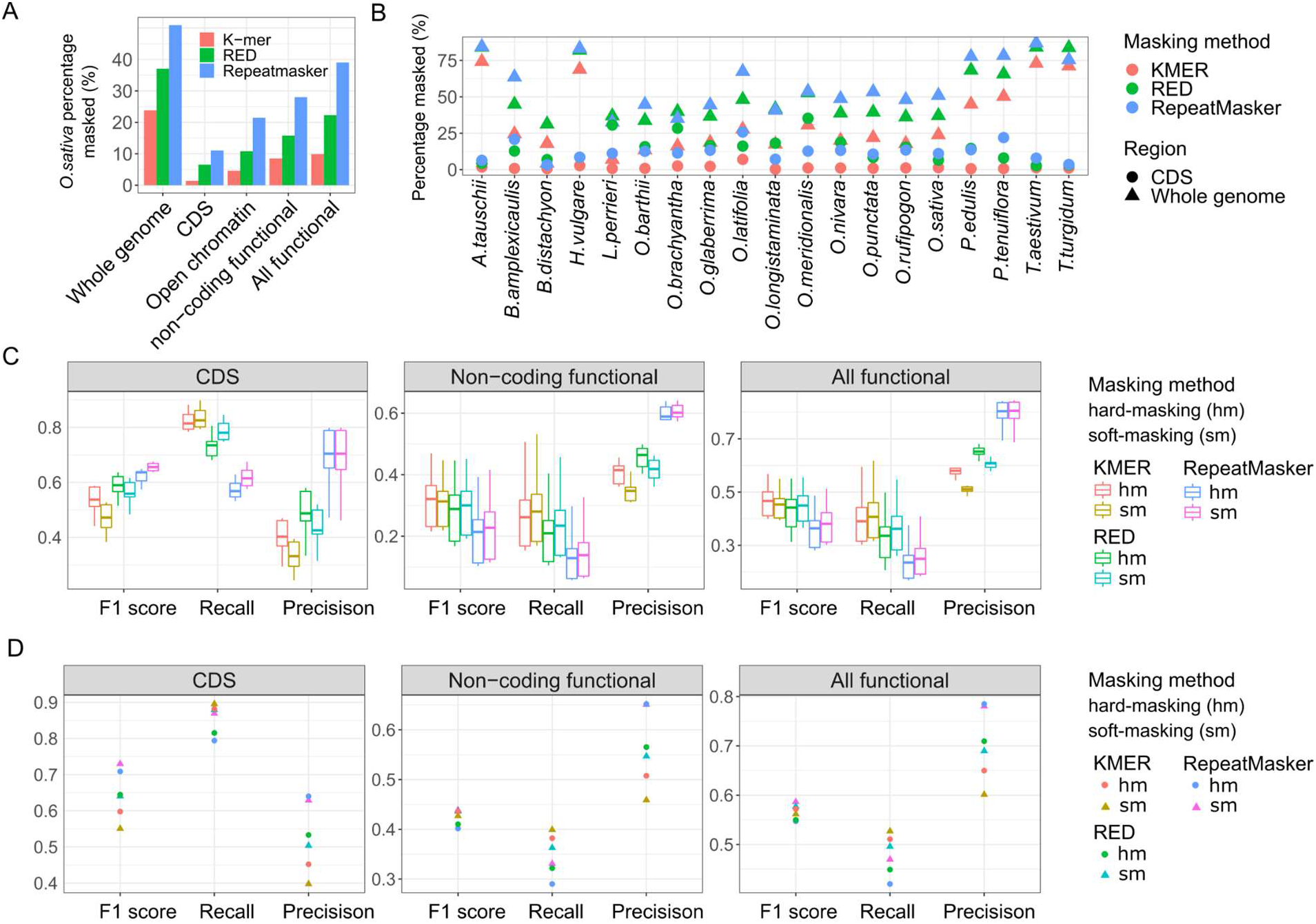
Impact of repeat masking methods on alignment of functional genomic regions in the BOP grass clade. (A) The masking rate for the whole genome, CDS, open chromatin, non-coding functional and all functional regions using three different masking methods in rice. (B) The masking rate for the whole genome (triangle) and for CDS regions (circle) in 19 species of the BOP clade. (C) Boxplots of pairwise alignment performance of each species in the BOP clade against the rice genome using different genome masking methods. (D) Multiple alignment performance in the BOP clade using different genome masking methods.

**Figure S2.**
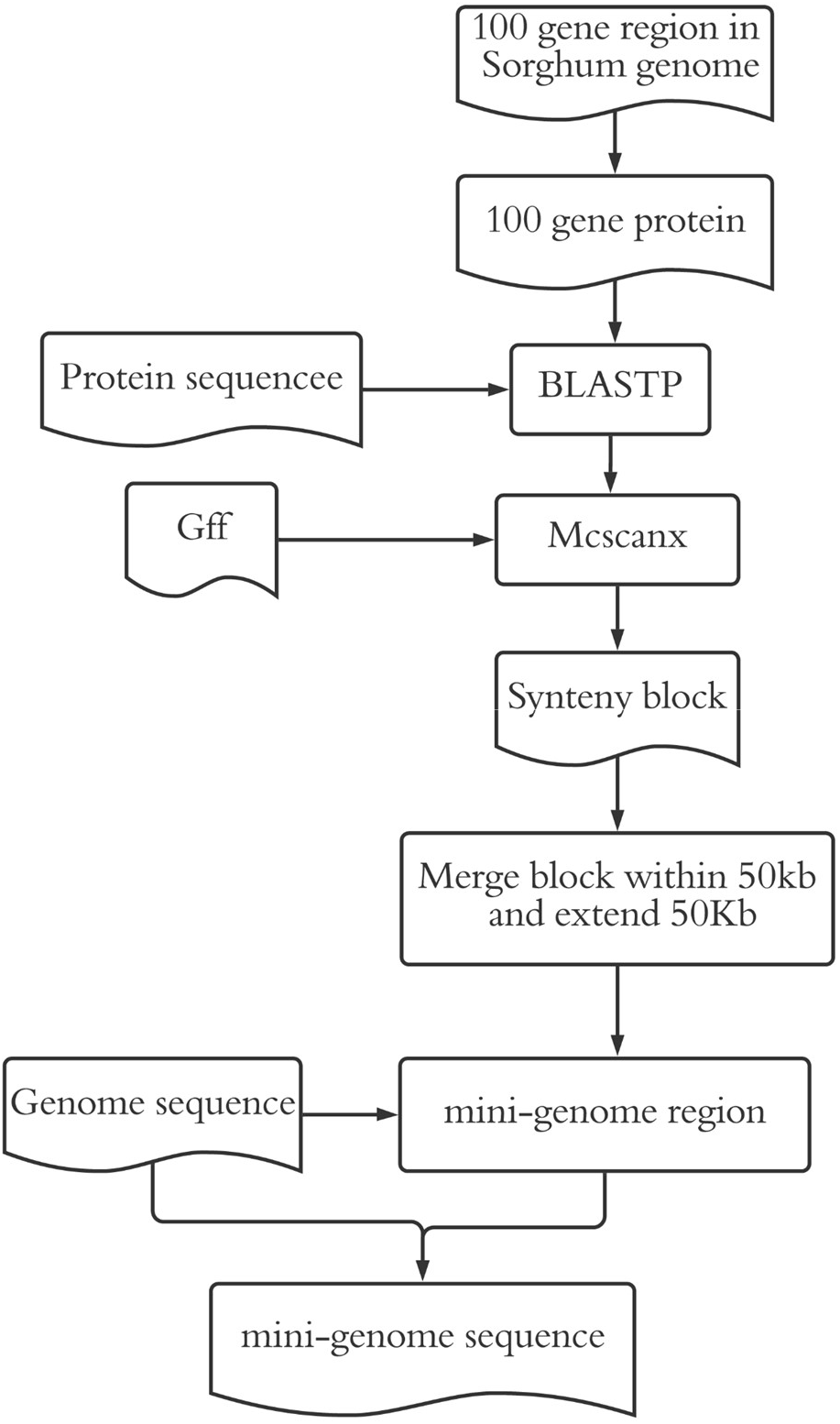
The mini-genome construction pipeline.

**Figure S3.**
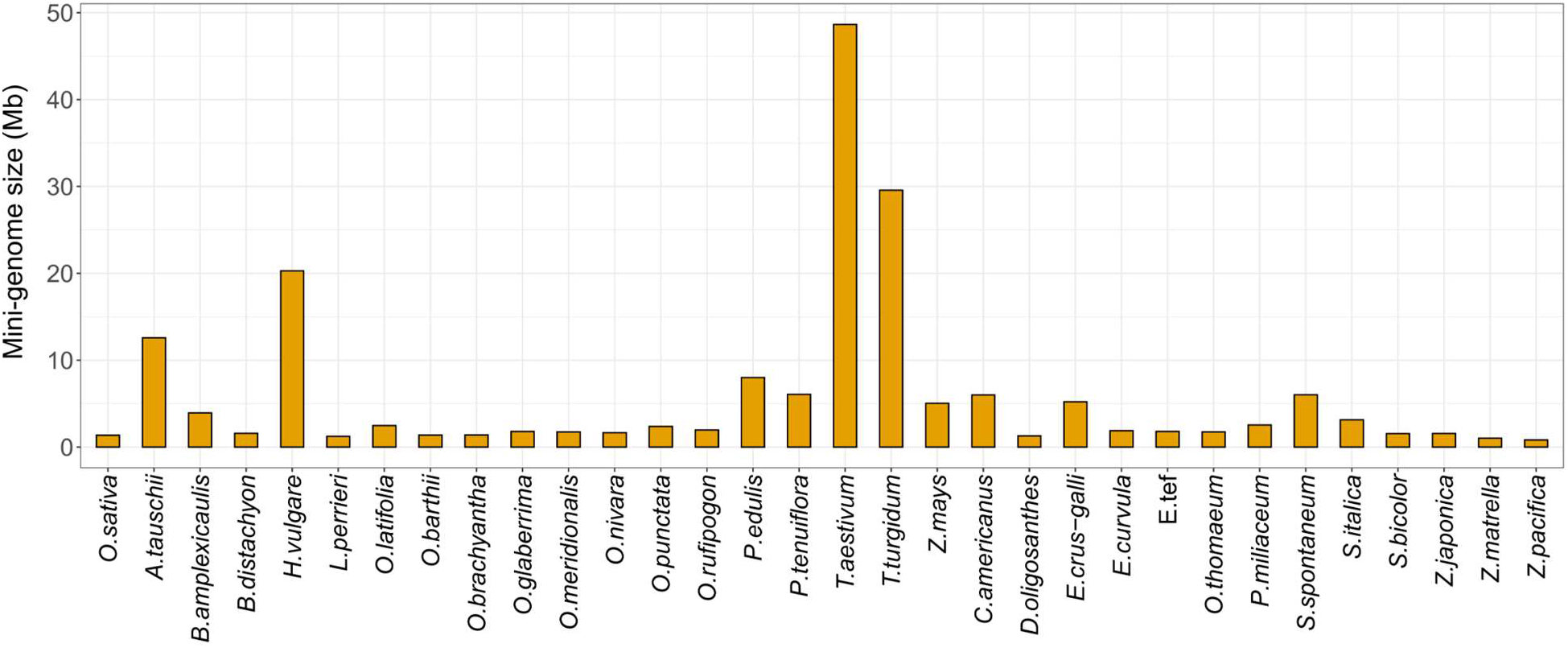
Comparison of mini-genome size.

**Figure S4.**
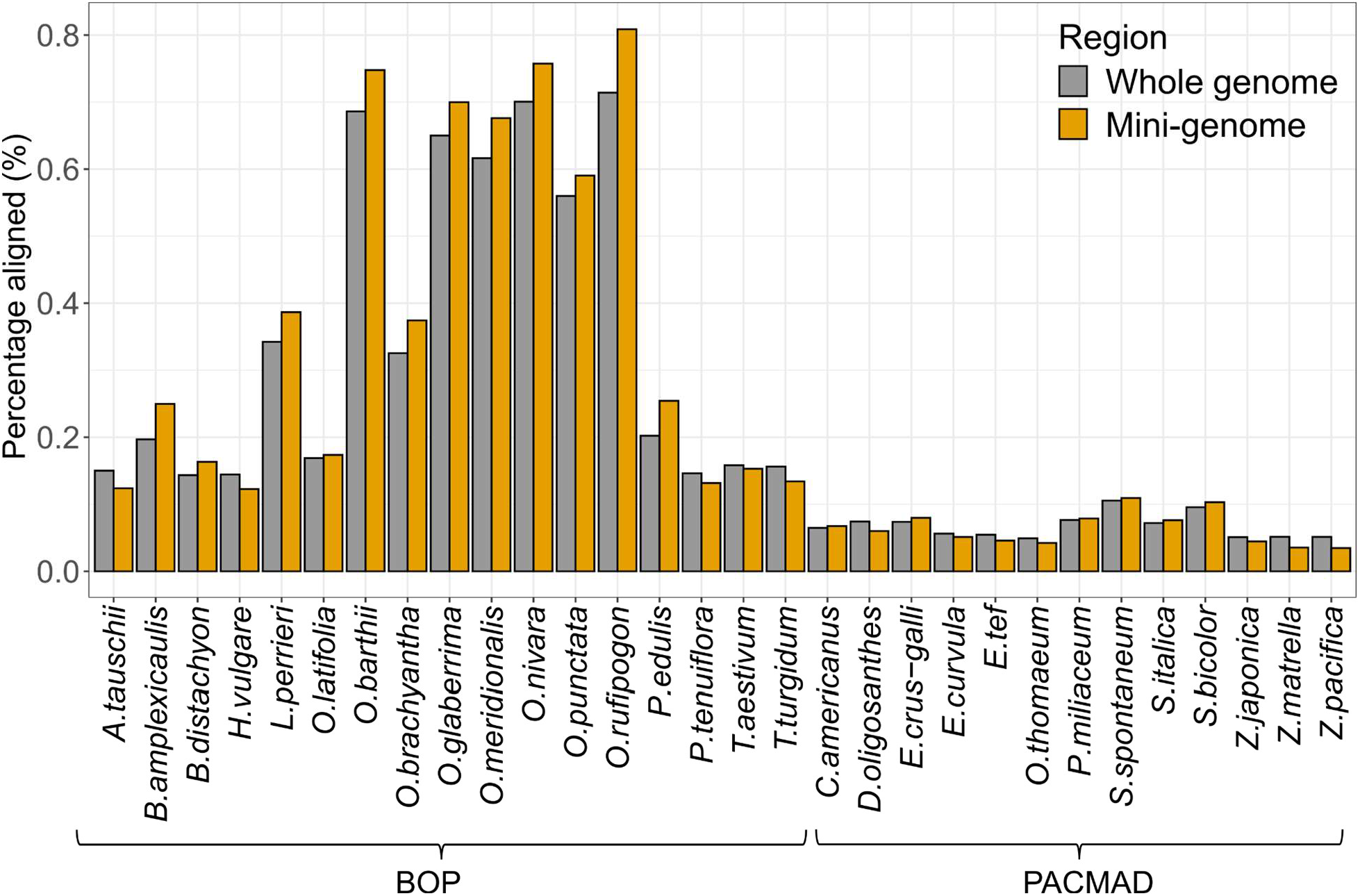
Comparison of the query-to-reference pairwise alignment rate for the whole genome and the mini-genome in 32 grass species. The grey bar is the alignment rate of each species aligned to reference species (*Zea mays* for PACMAD, *Oryza sativa* for BOP). The yellow bar is the alignment rate of each mini-genome aligned to reference the mini-genome.

**Figure S5.**
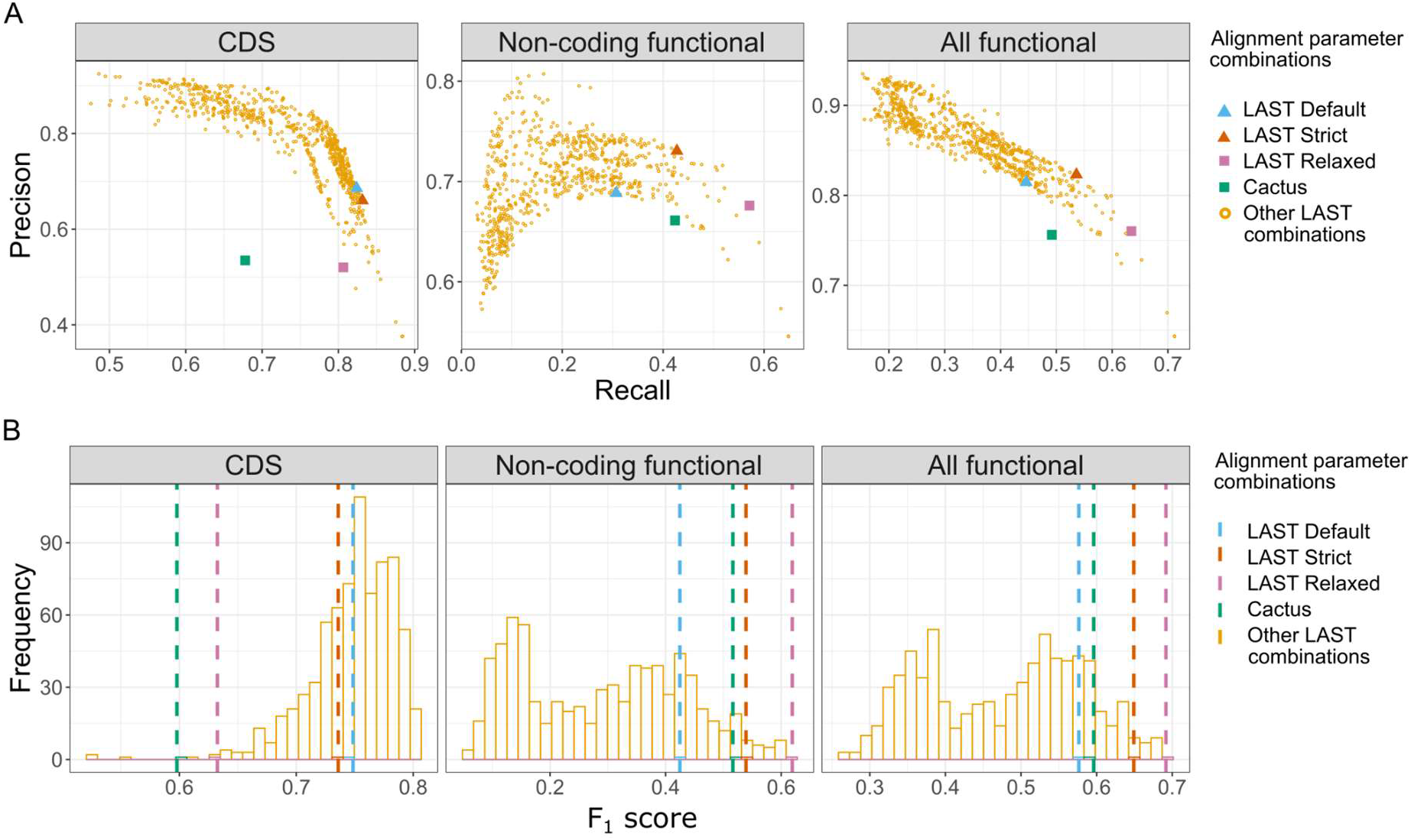
Multiple alignment performance of 750 LAST parameter combinations in different genomic regions in the BOP grass clade. Tested parameter combinations are compared to the alignment performance of default LAST parameters and the Cactus aligner based on (A) recall and precision as well as the (B) F_1_ score.

**Figure S6.**
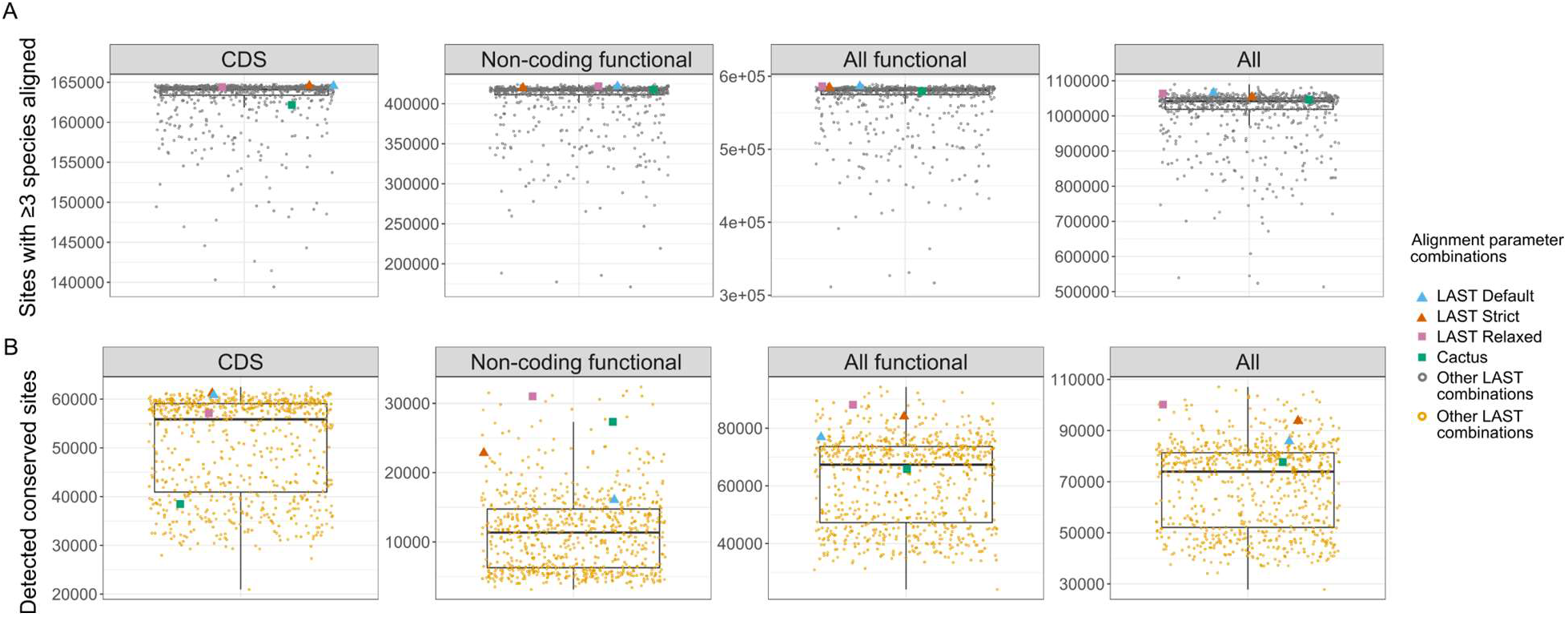
Conservation scoring based on multiple alignments generated with 750 different LAST parameter combinations for BOP clade.

**Figure S7.**
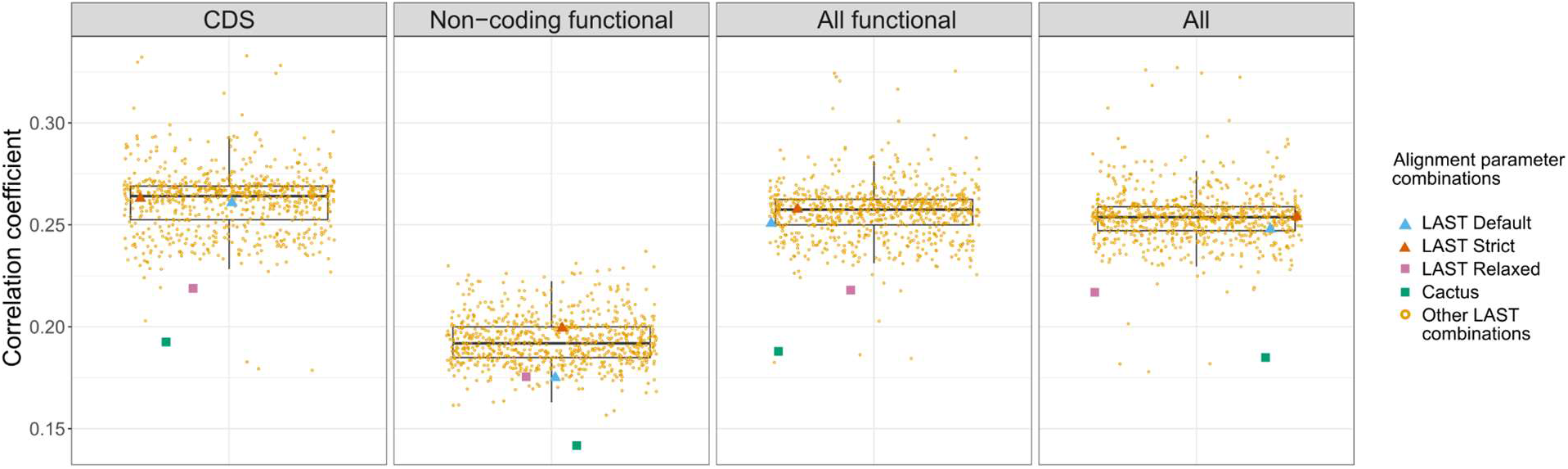
Correlation of GERP conservation scores between PACMAD and BOP clades for 750 LAST parameter combinations in different genomic regions.

**Supplementary Table S1.**
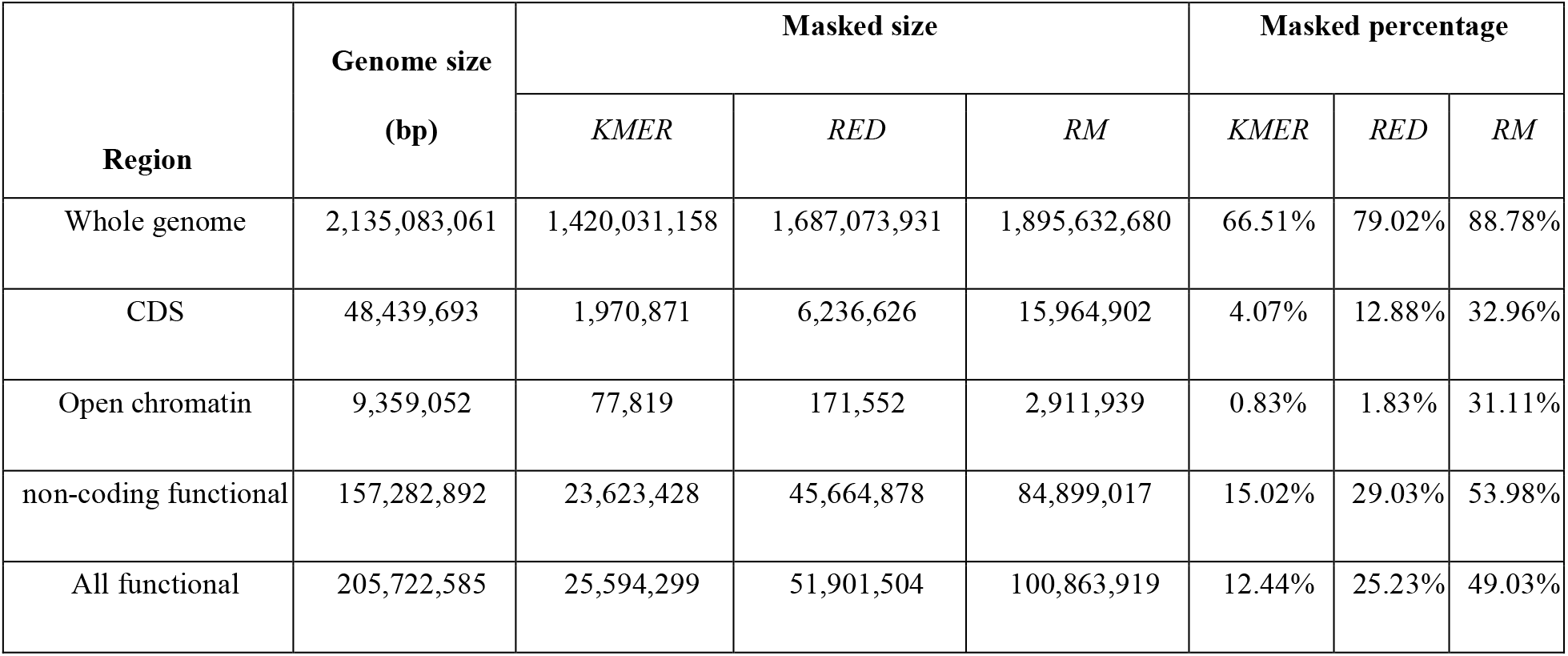
All three masking methods in *Zea mays* functional regions.

**Supplementary Table S2.**
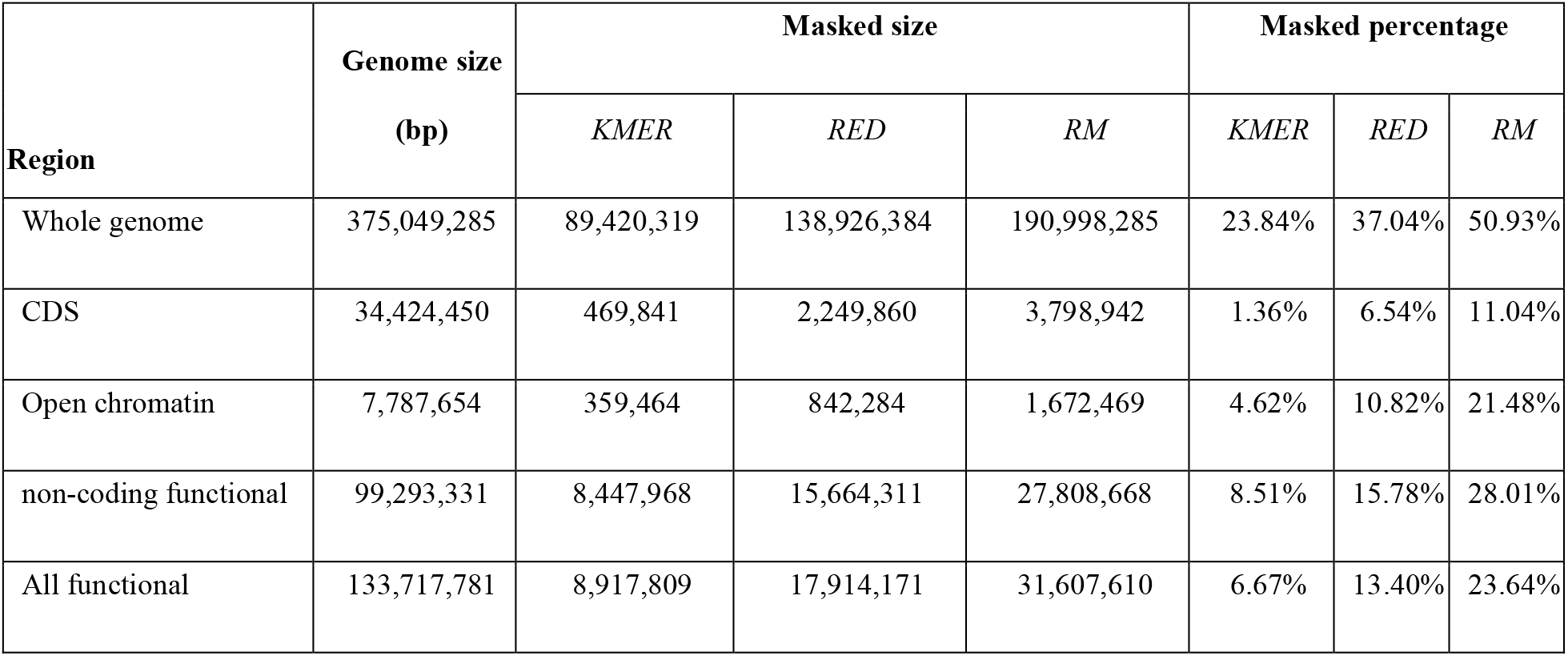
All three masking methods in *Oryza sativa* functional regions.

| Region | Genome size<br>(bp) | Masked size |  |  | Masked percentage |  |  |
| --- | --- | --- | --- | --- | --- | --- | --- |
|  |  | *KMER* | *RED* | *RM* | *KMER* | *RED* | *RM* |
| Whole genome | 375,049,285 | 89,420,319 | 138,926,384 | 190,998,285 | 23.84% | 37.04% | 50.93% |
| CDS | 34,424,450 | 469,841 | 2,249,860 | 3,798,942 | 1.36% | 6.54% | 11.04% |
| Open chromatin | 7,787,654 | 359,464 | 842,284 | 1,672,469 | 4.62% | 10.82% | 21.48% |
| non-coding functional | 99,293,331 | 8,447,968 | 15,664,311 | 27,808,668 | 8.51% | 15.78% | 28.01% |
| All functional | 133,717,781 | 8,917,809 | 17,914,171 | 31,607,610 | 6.67% | 13.40% | 23.64% |

**Supplementary Table S3.**
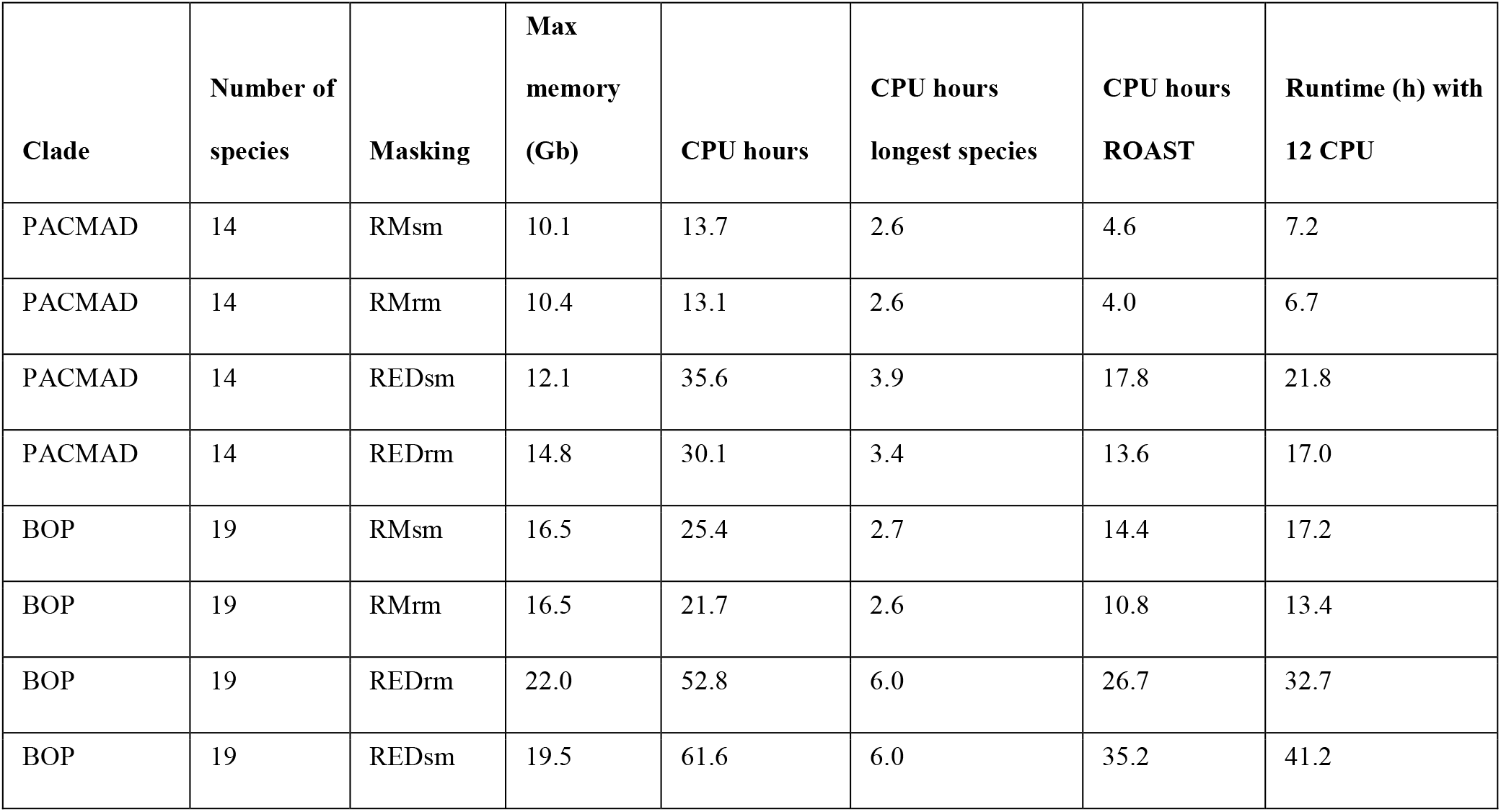
Memory usage and runtime for *msa_pipeline* using RepeatMasker (RM) and RED with soft-masking (sm) and hard-masking (rm).

| Clade | Number of species | Masking | Max memory (Gb) | CPU hours | CPU hours longest species | CPU hours ROAST | Runtime (h) with 12 CPU |
| --- | --- | --- | --- | --- | --- | --- | --- |
| PACMAD | 14 | RMsm | 10.1 | 13.7 | 2.6 | 4.6 | 7.2 |
| PACMAD | 14 | RMrm | 10.4 | 13.1 | 2.6 | 4.0 | 6.7 |
| PACMAD | 14 | REDsm | 12.1 | 35.6 | 3.9 | 17.8 | 21.8 |
| PACMAD | 14 | REDrm | 14.8 | 30.1 | 3.4 | 13.6 | 17.0 |
| BOP | 19 | RMsm | 16.5 | 25.4 | 2.7 | 14.4 | 17.2 |
| BOP | 19 | RMrm | 16.5 | 21.7 | 2.6 | 10.8 | 13.4 |
| BOP | 19 | REDrm | 22.0 | 52.8 | 6.0 | 26.7 | 32.7 |
| BOP | 19 | REDsm | 19.5 | 61.6 | 6.0 | 35.2 | 41.2 |

**Supplementary Table S4.** Alignment parameter ranges used for exploration of the parameter space. Step size of parameter value ranges shown in parentheses. See Supplementary Data for RETRO SIMPLE and RETRO custom matrices.

| *LAST*<br>Parameter | Default parameters for<br>two reference penalty<br>matrices |  | Parameter ranges for five penalty matrices |  |  |  |  |
| --- | --- | --- | --- | --- | --- | --- | --- |
|  | *HOXD70* | *1:1:1:7:1* | *1:1:1:7:1* | *2:1:2:16:1* | *RETRO SIMPLE* | *RETRO* | *HOXD70* |
| -R | 10 | 10 | 10,11 | 10,11 | 10,11 | 10,11 | 10,11 |
| -c | -c | - | -c | -c | -c | -c | -c |
| -u | MAM8 | YASS | YASS,MAM8 | YASS,MAM8 | YASS,MAM8 | YASS,MAM8 | YASS,MAM8 |
| -a | 400 | 7 | 3-16 (1) | 6-32 (2) | 3-16 (1) | 300-1600 (100) | 300-1600 (100) |
| -b | 30 | 1 | 1-3 (1) | 1-3 (1) | 1-3 (1) | 20-300 (10) | 20-300 (10) |
| -e | 4000 | 32 | 30-60 (10) | 60-120 (20) | 30-60 (10) | 3000-5000 (1000) | 3000-5000 (1000) |
| -m | 100 | 10 | 50-150 (50) | 50-150 (50) | 50-150 (50) | 50-150 (50) | 50-150 (50) |
| -y | - | 9 | 9-50 (5) | 15-60 (5) | 9-50 (5) | 1000-2000 (200) | 1000-2000 (200) |
| -x | - | 25 | 20-100 (10) | 30-150 (10) | 20-100 (10) | 1500-2500 (250) | 1500-2500 (250) |
| -d | - | 27 | e*0.7 | e*0.7 | e*0.7 | e*0.7 | e*0.7 |
| -p | HOXD70 | - | - | custom matrix | custom matrix | custom matrix | HOXD70 |
| -u | 2 | 2 | 1-3 (1) | 1-3 (1) | 1-3 (1) | 1-3 (1) | 1-3 (1) |

**Supplementary Table S5.**
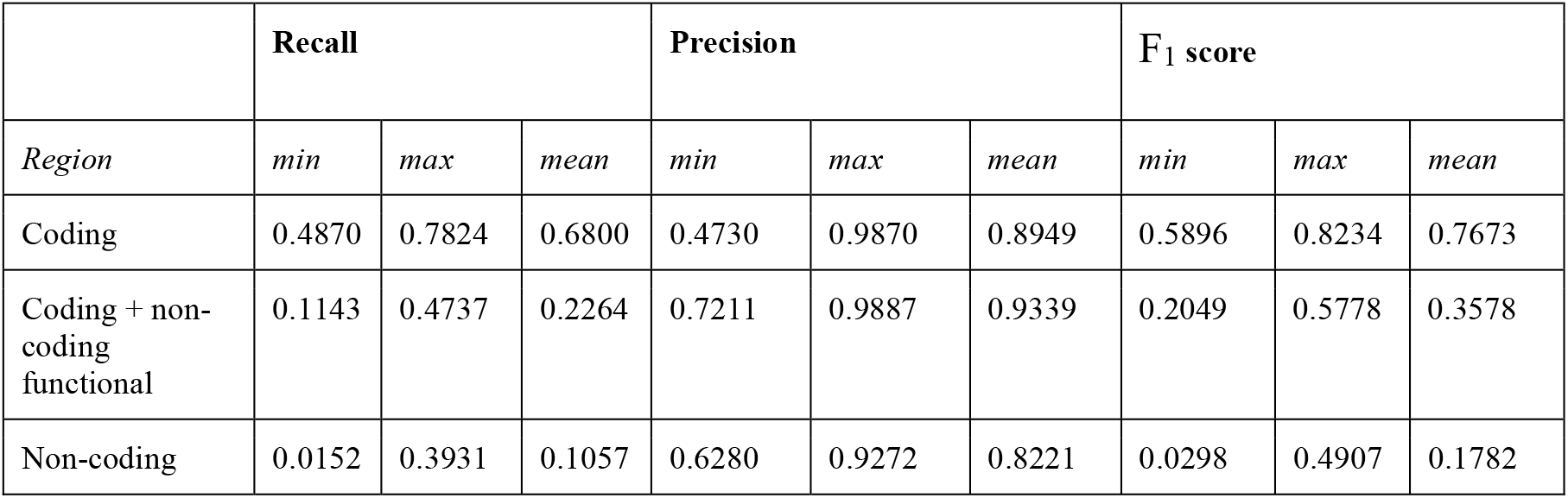
Alignment metrics in different genomic regions of the mini-genome based on 750 tested LAST parameter combinations.

**Supplementary Table S6.** Parameter settings of the LAST aligner at a default setting and at the relaxed and strict settings. The relaxed and strict settings were selected for divergent interspecies genome alignment from 750 parameter combinations tested in this study.

| Parameter | Parameter meaning | Default | Relaxed | Strict |
| --- | --- | --- | --- | --- |
| -R | repeat-marking options (10: do nothing, 11: carry out additional simple repeat masking with tantan) | 10 | 11 | 10 |
| -c | Exclude masked sequence from initial matches | no | yes | yes |
| -u | seeding scheme, sets matrix and sets penalty scores for lastal | YASS | YASS | YASS |
| -a | Gap/Insertion existence cost | 7 | 300 | 700 |
| -b | Gap extension cost | 1 | 20 | 20 |
| -e | Minimum alignment score | 32 | 5000 | 5000 |
| -m | Initial match number | 10 | 150 | 50 |
| -y | Maximum score drop for gapless alignments | 9 | 1800 | 1400 |
| -x | This option makes lastal extend gapped alignments twice. First, it extends gapped alignments with a maximum score drop of x | 25 | 1750 | 1750 |
| -d | Minimum score for gapless alignments. Can be set to equal -e value | 27 | 3500 | 3500 |
| -p | Mismatch matrix for alignment extension | equal mismatch weight matrix | HOXD70 | HOXD70 |
| -u | Specify treatment of lowercase letters when extending alignments (1:Mask them for gapless but not gapped extensions; 2: Mask them for gapless but not gapped extensions, and then discard alignments that lack any segment with score $\geq e$ when lowercase is masked) | 2 | 1 | 1 |

**Supplementary Table 7.** Mini-genome alignment metrics comparison for the Cactus aligner and three parameter combinations for the LAST aligner in the PACMAD clade.

|  | All functional region |  |  | Coding region |  |  | Non-coding functional region |  |  |
| --- | --- | --- | --- | --- | --- | --- | --- | --- | --- |
| Alignment | *F<sub>1</sub> score* | *recall* | *precision* | *F<sub>1</sub> score* | *recall* | *precision* | *F<sub>1</sub> score* | *recall* | *precision* |
| Cactus | 0.5472 | 0.4040 | 0.8478 | 0.7243 | 0.7637 | 0.6888 | 0.4404 | 0.3083 | 0.7706 |
| LAST default | 0.4309 | 0.2823 | 0.9095 | 0.7952 | 0.7483 | 0.8485 | 0.2652 | 0.1583 | 0.8166 |
| LAST strict | 0.5037 | 0.3483 | 0.9098 | 0.7825 | 0.7484 | 0.8199 | 0.3762 | 0.2418 | 0.8469 |
| LAST relaxed | 0.5685 | 0.4270 | 0.8500 | 0.7013 | 0.7337 | 0.6717 | 0.4795 | 0.3454 | 0.7836 |

**Supplementary Table S8.**
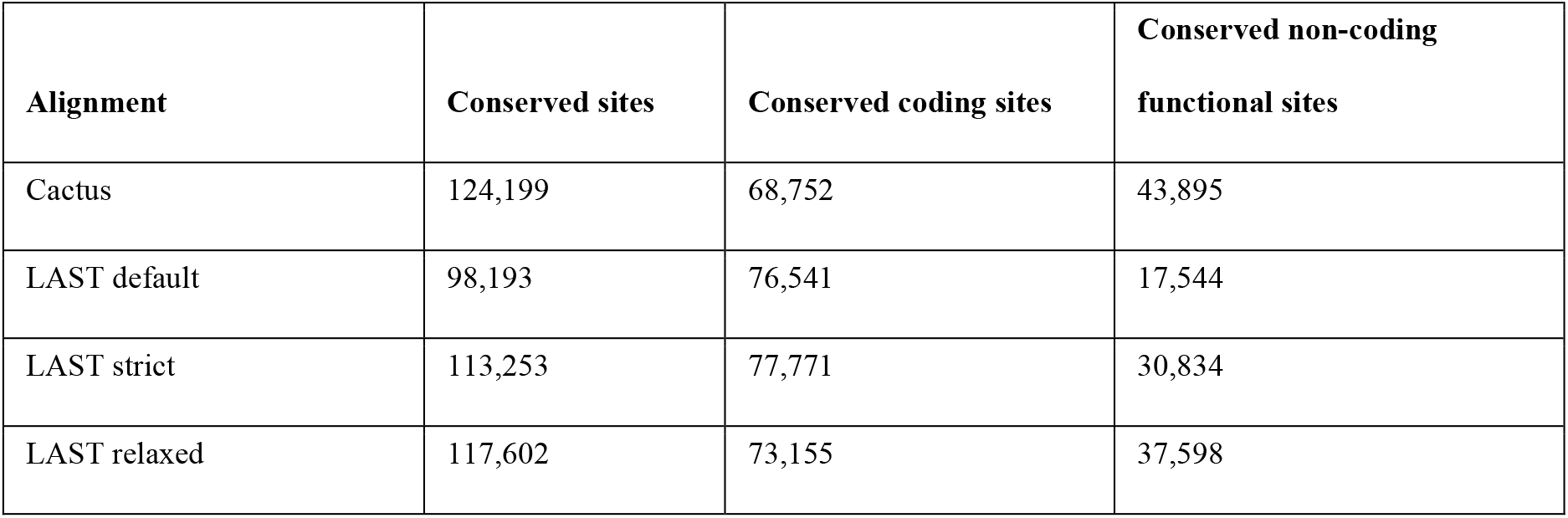
Conserved sites comparison in the PACMAD clade for the Cactus aligner and three parameter combinations for the LAST aligner.

